# Beyond performance: How design choices shape chemical language models

**DOI:** 10.1101/2025.05.23.655735

**Authors:** Inken Fender, Jannik Adrian Gut, Thomas Lemmin

**Affiliations:** Institute of Biochemistry and Molecular Medicine, University of Bern, Bühlstrasse 28, Bern, 3012, Bern, Switzerland; Graduate School for Cellular and Biomedical Sciences (GCB), University of Bern, Mittelstrasse 43, Bern, 3012, Bern, Switzerland

**Author notes:** Contributing authors. These authors contributed equally to this work (Inken Fender, Jannik Adrian Gut).

**Keywords:** large language models, chemical language models, interpretability, machine learning for chemistry, explainable AI (XAI), SMILES, SELFIES, RoBERTa, BART

## Abstract

Chemical language models (CLMs) have shown strong performance in molecular property prediction and generation tasks. However, the impact of design choices, such as molecular representation format, tokenization strategy, and model architecture, on both performance and chemical interpretability remains underexplored. In this study, we systematically evaluate how these factors influence CLM performance and chemical understanding. We evaluated models through finetuning on downstream tasks and probing the structure of their latent spaces using simple classifiers and dimensionality reduction techniques. Despite similar performance on downstream tasks across model configurations, we observed substantial differences in the structure and interpretability of their internal representations. SMILES molecular representation format with atomwise tokenization strategy consistently produced more chemically meaningful embeddings, while models based on BART and RoBERTa architectures yielded comparably interpretable representations. These findings highlight that design choices meaningfully shape how chemical information is represented, even when external metrics appear unchanged. This insight can inform future model development, encouraging more chemically grounded and interpretable CLMs.

**Scientific Contribution:** This study systematically evaluates how core design choices influence chemical language models. Although the performances on downstream tasks were often similar across configurations, we observed substantial differences in internal representations with atomwise tokenized SMILES representations producing more chemically structured latent spaces than representations based on SELFIES. By clarifying the effects of molecular representation format and tokenization strategy, our findings provide actionable guidance for the more informed and interpretable design of future CLMs.

**Graphical Abstract:** 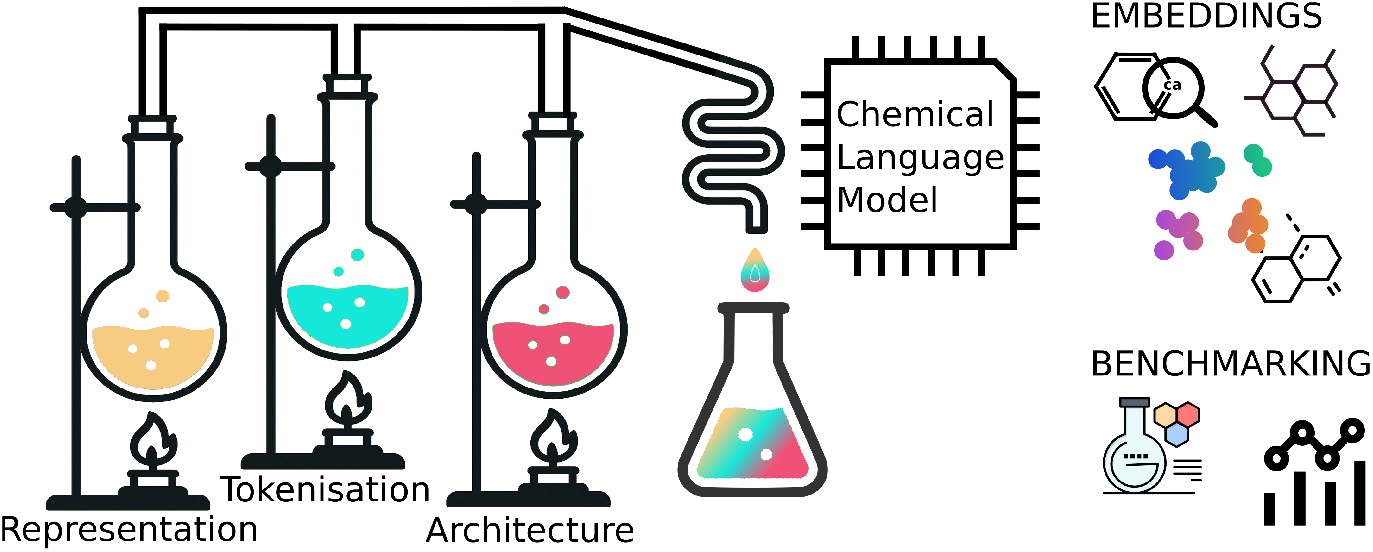

## 1 Introduction

The design and discovery of novel molecules with desired properties are crucial for advances in medicine, materials science, and agriculture. Traditional experimental methods are often time-consuming and expensive, driving the need for efficient computational approaches. As a result, computational methods have become essential tools for accelerating molecular innovation.

Recent advances in deep learning, particularly in natural language processing (NLP), have sparked growing interest in applying language models to molecular data. By representing molecules as sequences, most commonly using SMILES (Simplified Molecular Input Line Entry System) [1] strings, text-based cheminformatics models have demonstrated strong performance across a range of tasks, including prediction of molecular properties, completion of reactions, planning of retrosynthesis, and generation of de novo molecules [2–5]. In several cases, these models have even outperformed earlier cheminformatics methods such as molecular fingerprint-based methods or GNNs [4, 6, 7].

To improve the syntactic robustness of sequence-based molecular representations, SELFIES (Self-Referencing Embedded Strings) [8] was introduced as an alternative to SMILES. Unlike SMILES, which can produce invalid molecules due to syntax errors, every SELFIES string maps to a valid molecule by design. In addition, tokenization strategies play a crucial role in how models interpret molecular sequences. The most common approaches include atomwise tokenization [4], which segments the string based on atoms and bonds, and subword tokenization using SentencePiece [9, 10], which learns data-driven tokens optimized for training efficiency on a finer level.

Ultimately, the performance and interpretability of these models are influenced by several design choices, including molecular representation, tokenization strategies and the architectural backbone. These factors collectively influence both the downstream task performance and the internal representation of the chemical knowledge of the model. However, a complete understanding of the individual and synergistic effects of these components is still lacking, motivating increased research in this area [11, 12]. Although some initial evidence suggests that the choice of molecular representation might not be a primary driver of property prediction performance [2, 13, 14], the impact of tokenization [12] and, crucially, the architectural backbone [15] are still poorly understood and require more in-depth investigation. While non-canonical SMILES have shown benefits in certain settings, prior work recommends canonization when computational resources are limited, as it improves training efficiency without sacrificing performance [16].

In this work, we therefore systematically investigate the impact of three core design choices in large chemical language models (CLMs): molecular representation (SMILES vs. SELFIES), tokenization strategy (atomwise vs. SentencePiece), and model architecture (RoBERTa vs. BART). Our goal is to understand how these choices influence downstream performance, the structure of the latent space, and the chemical interpretability of the learned embeddings. We evaluate models on predictive tasks, probe the organization of their latent representations using simple classifiers and dimensionality reduction, and examine atom-level embeddings in relation to chemical typing schemes. While downstream task performance is often comparable across configurations, we find that certain setups, particularly those using SMILES with atomwise tokenization, yield more chemically structured embeddings, potentially indicating a deeper internalization of chemical context.

## 2 Methods

We investigated the performance of Transformer-based models for chemical structure representation. We trained a series of BART and RoBERTa models, exploring the impact of varying tokenization strategies (atomwise and SentencePiece) and molecular representations (SMILES and SELFIES).

### 2.1 Pretraining datasets

The initial dataset was derived from the PubChem-10M dataset [3], a collection of 10 million chemical structures sourced from PubChem [17]. To ensure the quality and consistency of the SMILES strings, all molecules were canonicalized using the RDKit [18]. For each molecule, the RDKit was employed to generate potential isomers, with a maximum of ten attempts per molecule. These isomers allow to explicitly model chirality. Subsequently, to generate the corresponding SELFIES representations, the canonicalized SMILES were converted to SELFIES using the SELFIES library [8]. To validate the accuracy of the SELFIES conversion and ensure reversibility, the generated SELFIES strings were then back-translated to SMILES using the same library. Only molecules for which the back-translated SMILES matched the original canonicalized SMILES were retained. Due to this procedure, 10’818 (0.1%) out of the ten million molecules in the base dataset were filtered out and 5’460’790 isomers were added with explicit chirality.

### 2.2 Tokenization

Two different tokenization strategies were used: atomwise [4] and SentencePiece [9, 10]. The atomwise strategy decomposes SMILES strings into individual atoms and bonds, treating each as a separate token. Alternatively, SentencePiece utilizes a subword tokenization technique, grouping characters within and across atoms based on their frequency of occurrence. To generate the SentencePiece vocabulary, we utilized the Hugging Face Transformers library [19], creating a vocabulary of 1’000 subword units.

### 2.3 Language model description

Two Transformer-based large language model architectures were employed: BART [20] and RoBERTa [21]. Both models leverage the Transformer architecture [22] to capture long-range dependencies within sequential data. BART (Bidirectional and Auto-Regressive Transformers) is an encoder-decoder model designed for sequence-to-sequence tasks. It utilizes a denoising autoencoder objective during pretraining, where corrupted input sequences are reconstructed. RoBERTa (Robustly Optimized BERT Pretraining Approach) is an encoder-only model that builds upon the BERT [23] architecture. It is pretrained using a masked language modelling objective, where the model predicts masked tokens within input sequences.

Both BART and RoBERTa models were implemented and trained using the fairseq library [24]. Detailed training parameters, including hyperparameters and optimization settings, are provided in Section S1 in the Supplementary Information.

### 2.4 Downstream tasks

To evaluate the performance of our pretrained models, we conducted fine-tuning experiments on a series of downstream tasks sourced from the MoleculeNet benchmark suite [25, 26]. The following classification tasks were used: BACE, BBBP, ClinTox (CT TOX), HIV, and Tox21 (SR-p53). For regression tasks, we utilized the BACE, Clearance, Delaney, and Lipo tasks. We employed the default train-validation-test splits provided by MoleculeNet for each dataset. The data processing pipeline mirrored that of the pretraining dataset, with the exception of explicit isomer generation. We note that there are eight samples (0.18%) in the HIV test set that did not pass the SELFIES filters and are omitted in all our tests.

### 2.5 Atom type embedding analysis

To evaluate the ability of the models to cluster and distinguish different atom types, we created mol2-files for 973 molecules from the BBBP, Clearance, Delaney, BACE classification and Lipo datasets and attempted to assign GAFF2 atom types to each atom of those molecules using Antechamber [27]. To assign atom types to SELFIES, the SMILES were translated to SELFIES using the selfies library with the flag “attribute=True” to track what SMILES token led to the creation of which SELFIES token. This ensured consistent atom type assignment across both input formats.

To ensure the reliability of the atom type assignments, we used parmchk [27] to filter out molecules with atom assignments that resulted in a penalty score exceeding 300, indicating potential inconsistencies or errors in the parametrization.

For each molecule, we then extracted the embedding vector corresponding to each atom from our pretrained models. To facilitate a fair comparison, we only analysed embeddings for atom types that were consistently assigned across both SMILES and SELFIES representations. This led us to a final analysis of 143 molecules with atom assignments and embeddings across different models and representations. Principal Component Analysis (PCA) was subsequently used to visualize the distribution and similarity of these atom type embeddings in the embedding space. The maximum number of depicted atoms per type was limited to 300.

## 3 Results

### 3.1 Downstream task performance

We first pretrained 16 distinct models, systematically varying key architectural and data representation parameters. Specifically, we explored combinations of two molecular representations (SMILES [1], SELFIES [8]), two model architectures (BART [20], RoBERTa [21]), two tokenization strategies (atomwise, SentencePiece [9]), and two chirality representations (implicit, explicit).

To evaluate the impact of these parameters on the models’ predictive capabilities, we performed fine-tuning experiments on a series of downstream tasks from the MoleculeNet datasets [25] of the DeepChem benchmark suite [26]. Each of the 16 pre-trained models was fine-tuned on five classification tasks (BACE, BBBP, ClinTox, HIV, Tox21) and four regression tasks (BACE, Clearance, Delaney, Lipo) and after hyperparameter tuning, fine-tuning was repeated five times using different random seeds to assess model robustness.

Given the varying difficulty of these downstream tasks, direct comparison of raw performance metrics is challenging. Therefore, we present and discuss the z-scores [28] of the models’ performance (Figure 2), which were computed to normalize the performance of each model across the different tasks. For classification tasks, z-scores were calculated based on the area under the precision-recall curve (PR-AUC). For regression tasks, z-scores were calculated for each rectified root mean squared error (RMSE) and then averaged. Raw performance metrics and comparisons to benchmark models for classification and regression tasks are available in Supplementary Information Table S3 and Table S4 respectively.

**Fig. 1.**
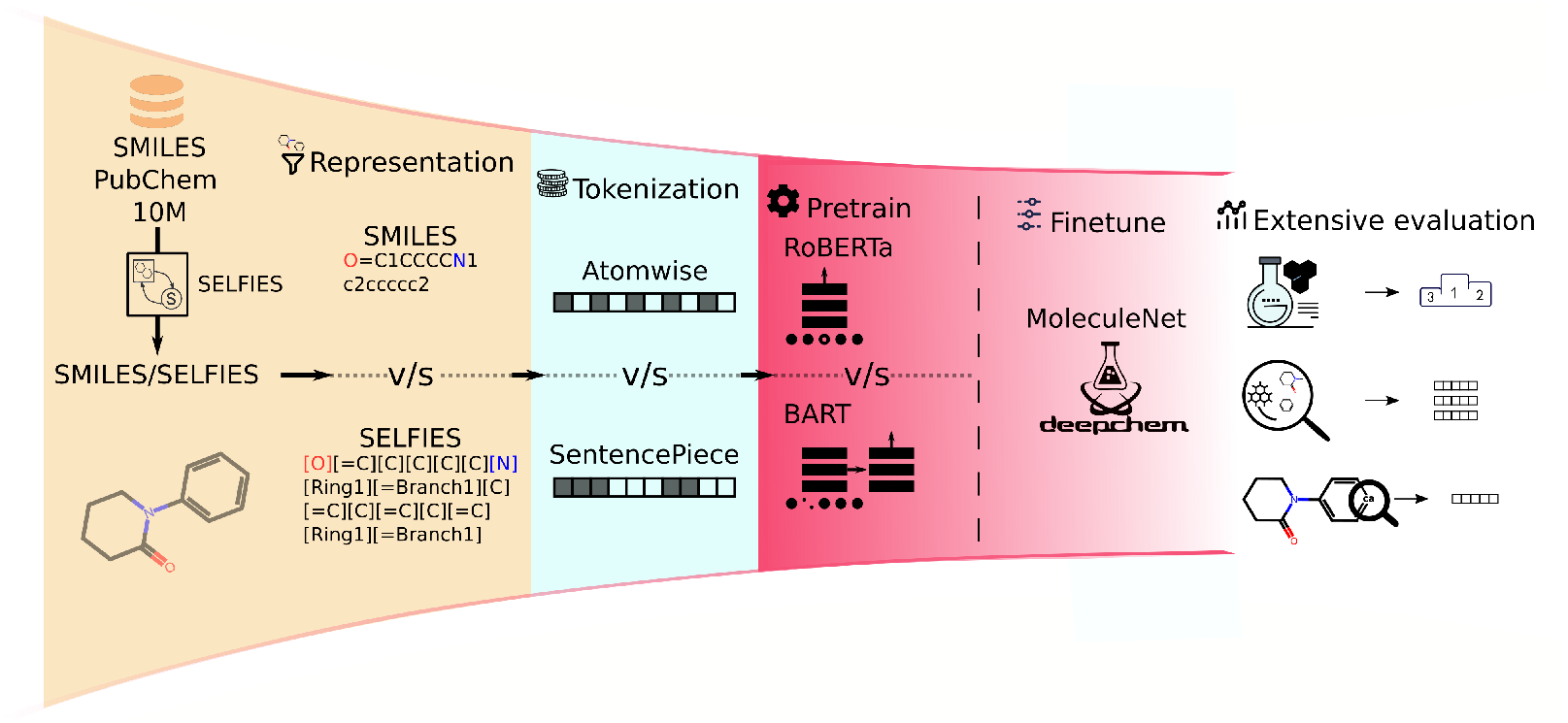
Workflow of the systematic analysis of molecular representation, tokenization strategies, and architecture in chemical language models (CLMs).

**Fig. 2.**
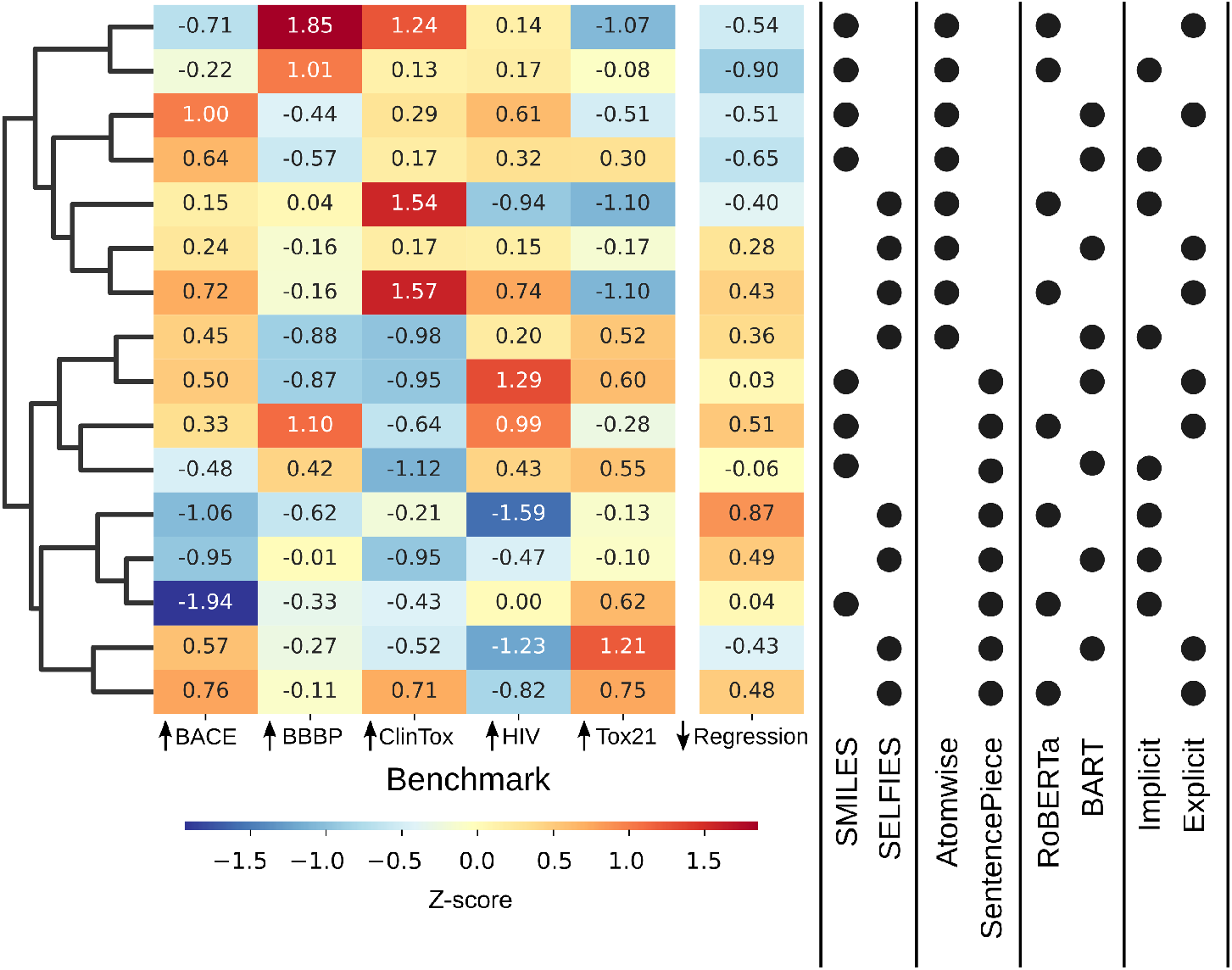
Z-scores of different tasks of different model configurations grouped by their cosine similarity. Z-scores were calculated from PR-AUC, except for the aggregated regression score which is based on RMSE. Model configurations are listed, starting from left to right, by molecule embedding, tokenisation, language model architecture and chirality representation.

While no single model configuration consistently outperformed all others, hierarchical clustering of models, based on the cosine similarity of their z-score performance profiles, revealed meaningful groupings. The first-level clustering mainly separated models by tokenization strategy. Notably, models employing atomwise tokenization generally exhibited superior performance compared to those using SentencePiece, with the exception of the Tox21 task.

At the next level of clustering, models further grouped according to the molecular representation used. SMILES representations tended to yield better performance on the BBBP and HIV tasks as well as aggregated regression scores, whereas SELFIES representations showed advantages on the ClinTox task. Explicit chirality representation was associated with improved performance on the BACE, ClinTox, HIV, Tox21, and BBBP tasks.

The model architecture influenced performance. BART models demonstrated better results on the BACE, HIV, and Tox21 tasks, while RoBERTa models achieved higher performances on the ClinTox and BBBP task.

Based on Wilcoxon signed-rank tests [29] (Table S5 in the Supplementary Information), we prioritized atomwise tokenization due to its statistically significant superior performance (p = 0.020) and implicit chirality representation, which showed no significant detriment (p = 0.123). For subsequent analyses, we further investigated BART and RoBERTa architectures with both SMILES and SELFIES.

### 3.2 Latent space analysis

To assess whether our pretrained models captured meaningful chemical features within their latent spaces, we investigated the ability of these spaces to encode two non-trivial molecular properties: the presence of heterocycles and the presence of at least one hydrogen bond donor.

For this analysis, we sampled 100’000 molecules for each feature class (presence/absence) from our pretraining dataset. These molecules were annotated with the target features using the RDKit [18]. Subsequently, the latent embeddings of each molecule were generated with pretrained models. These embeddings were then randomly split into equal-sized training and test sets.

We trained and evaluated three simple classifiers [30] on these latent embeddings to probe the structure of the latent space (details in Section S2 of the Supplementary Information). Each classifier employs a distinct decision boundary, revealing different aspects of how molecules are organized based on the target features. Specifically, k-Nearest Neighbors (k-NN) [31] assesses local density, classifying molecules based on the majority class among their nearest neighbours, providing insight into local clustering patterns. The Support Vector Classifier (SVC) with Radial Basis Function (RBF) kernel [31, 32] identifies complex, non-linear relationships, revealing if the latent space allows for intricate separation of molecules based on the target features. Finally, LinearSVC [33] tests global linear separability, determining if a simple hyperplane can effectively distinguish molecules based on the presence or absence of features.

BART embeddings consistently surpassed RoBERTa across all classification tasks, with the most pronounced advantage observed in the LinearSVC tests (Figure 3). Within BART embeddings, SMILES representations slightly outperformed SELFIES. In contrast, RoBERTa embeddings showed comparable performance between SMILES and SELFIES, except for a substantial performance deficit of SMILES in LinearSVC for heterocycle prediction.

**Fig. 3.**
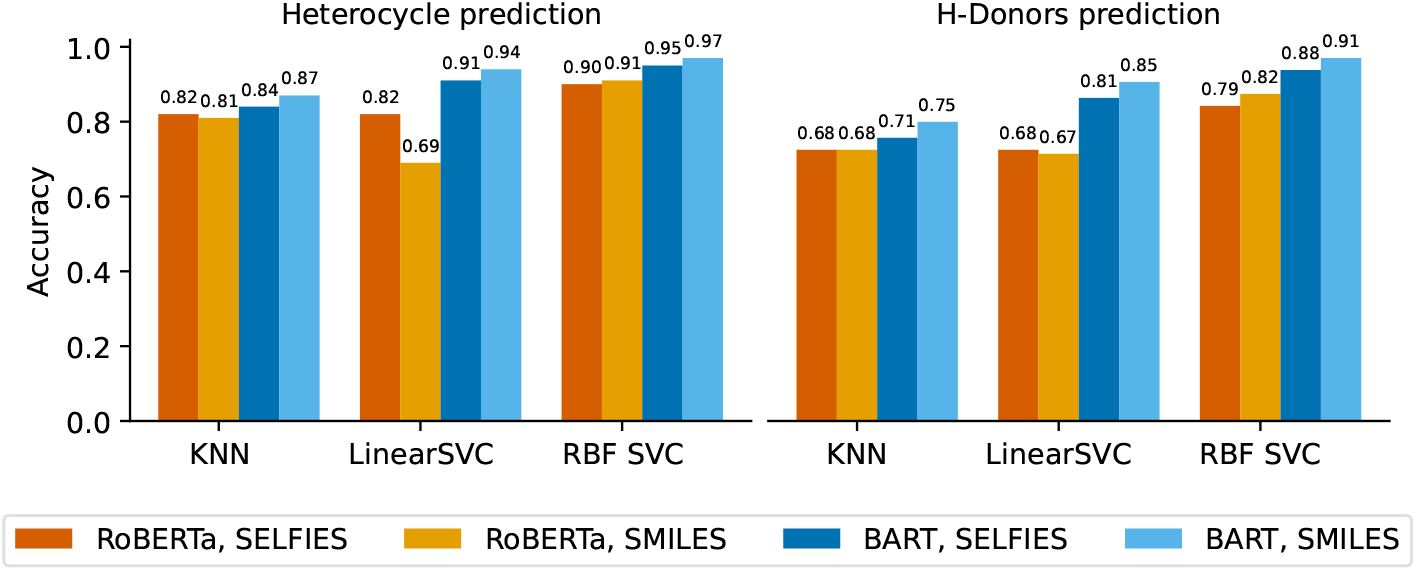
Accuracy of three simpler scikit-learn classifiers on heterocycle and H-donor prediction of four pretrained models with different language model architectures and molecule representation with the atomwise tokeniser and implicit chirality representation.

The RBF-SVC consistently yielded the highest performance across all models and tasks, suggesting the presence of complex, non-linear relationships in the latent space that effectively separate molecules based on target features. The choice between LinearSVC and k-NN as the second-best classifier depends on the underlying embedding model. For BART, LinearSVC exhibited superior performance, whereas for RoBERTa, k-NN and LinearSVC performed comparably.

These results suggest that local molecular neighbourhoods, as captured by k-NN, are similarly indicative of heterocycle presence and H-donor capabilities, regardless of the embedding model or molecular representation (SMILES/SELFIES). However, the SVC tests reveal that latent space of BART offers clearer, more distinct boundaries for these properties than RoBERTa. The difference in performance between LinearSVC and RBF-SVC, reflecting the degree of non-linearity required for optimal classification, was relatively small for BART-based models, indicating a more linearly separable latent space, but larger for RoBERTa, highlighting the need for more complex, non-linear decision boundaries.

### 3.3 Molecule embeddings

For a qualitative assessment of the learned embedding space, we selected 64 molecules from each of the following four chemical classes: steroids, beta-lactams, tropanes, and sulfonamides. These chemical classes were chosen because they all have some common use in pharmacology and yet have distinct structures and chemical features. The selected molecules were embedded using the atomwise tokenizer with implicit chirality representation.

BART-based SMILES embeddings exhibited the most distinct clustering in the UMAP [34] plot (Figure 4), with well-defined chemical families showing different degrees of separation. In contrast, BART-based SELFIES embeddings also demonstrated structured clustering, but the separation was slightly less pronounced, although beta-lactam clusters remained clearly identifiable. RoBERTa-based SMILES embeddings showed less definition than their BART counterparts, whereas RoBERTa-based SELFIES embeddings produced a more scattered distribution, with only weak clustering tendencies.

**Fig. 4.**
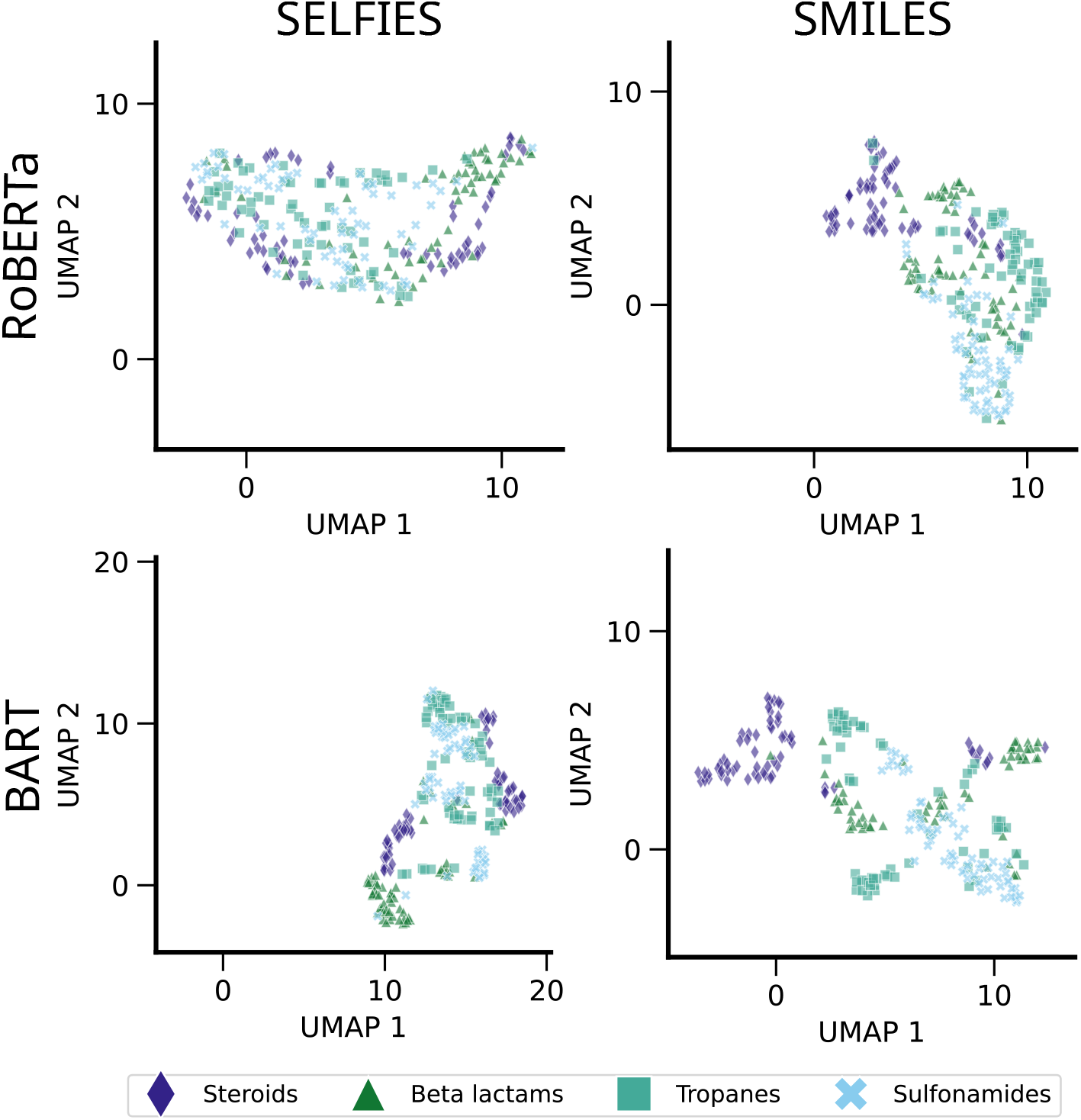
UMAP visualization of molecular embeddings; Scatter plot depicting the latent space organization of 64 molecules from four distinct chemical families each: steroids (purple diamonds), beta-lactams (green triangles), tropanes (teal squares), and sulfonamides (blue crosses). Embeddings were generated from SELFIES or SMILES representations using RoBERTa or BART models with atomwise tokenization and implicit chirality representation.

To further validate these observations, we performed a complementary PCA analysis of the selected molecules (Figure S1 in the Supplementary Information). For BART with SMILES, the first principal component was sufficient to separate sulfonamides and steroids, although beta-lactams and tropanes still formed a combined cluster. BART with SELFIES required two principal components to distinguish beta-lactams and steroids, while sulfonamides exhibited only a weak clustering pattern. The PCA results for RoBERTa revealed that its first principal component explains a substantially larger portion of the variance compared to BART, although both models are trained using standard hyperparameters (Section S1 in the Supplementary Information). The higher explained variance captured by the first principal component in RoBERTa compared to BART persists across all combinations of other design choices, though the difference is less pronounced (Figure S1 in Supplementary Information). RoBERTa with SMILES effectively identifies steroids using the first principal component, while the second component helps to identify sulfonamides. In contrast, RoBERTa with SELFIES only shows weak clustering tendencies without clear separation.

To ensure that the observed clustering patterns arise from meaningful learned features rather than artifacts of the embedding process or dimensionality reduction techniques, we evaluated an untrained model as a control (Figure S2 in the Supplementary Information). In the untrained PCA projection, sulfonamides form a distinct cluster, likely due to the presence of sulphur, a feature absent in the other molecules, making it identifiable even without training. However, the remaining molecular classes do not exhibit clear separation, with their embeddings appearing mixed. Similarly, the UMAP projection shows weak clustering tendencies, but no well-defined clusters emerge. These results suggest that the structured organization observed in the learned embedding of the trained models captures meaningful chemical information rather than reflecting artifacts of the embedding or dimensionality reduction techniques.

### 3.4 Atom type embeddings

To evaluate the degree of chemical understanding acquired by our pretrained models, we analysed the similarity of atom embeddings based on GAFF2 atom types [27]. The goal was to determine whether atoms sharing the same GAFF2 atom type are represented by similar embedding vectors and whether their clustering differs between the different models. For this analysis, we focussed on the most commonly occurring atom types.

For carbon atoms, PCA of the embeddings reveals distinct clustering patterns across molecular representations and architectures (Figure 5). In the RoBERTa model trained on SMILES, aromatic carbons (GAFF2 atom type: ca) are effectively separated from sp^2^-hybridized aliphatic or ketone/thioketone carbons (GAFF2: c2 and c) and sp^3^ carbons (GAFF2: c3). A similar, albeit not as strong separation can be seen for BART trained on SMILES as well. When aromatic carbons are excluded and the PCA is rerun, the RoBERTa SMILES model still distinguishes ketone/thioketone carbons from sp^3^ carbons along the first principal component, with a similar separation observable along the second component in the RoBERTa SELFIES model (Figure S5 in the Supplementary Information). The BART models, regardless of whether they use SMILES or SELFIES, show less clear separation (Figure S5 in the Supplementary Information).

**Fig. 5.**
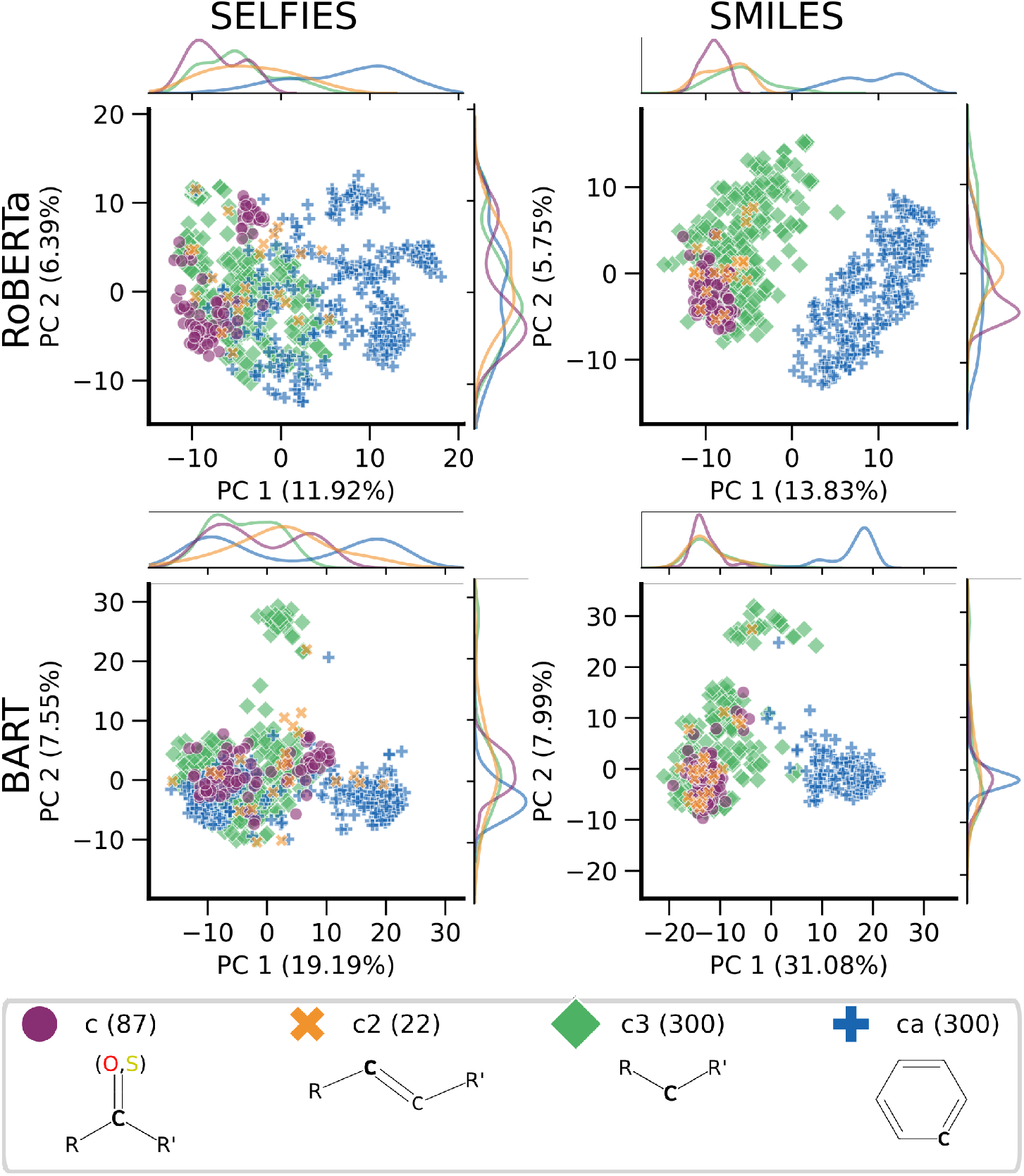
PCA of carbon atom type embeddings of SMILES or SELFIES-based models BART and RoBERTa with atomwise tokeniser and implicit chirality. The GAFF2 atom types have been determined by antechamber[27] and correspond to the following hybridizations: c: sp^2^ in C=O, C=S, c2: sp^2^ in aliphatic carbon, c3: sp^3^, ca: sp^2^ in aromatic carbon. Amount of examples in brackets.

For nitrogen atoms, the embeddings reveal that the sp^2^-hybridized nitrogen in aromatic rings (GAFF2: na) clusters distinctly in the RoBERTa SMILES model along the first principal component (Figure S3 in the Supplementary Information). In this model, sp^2^-hybridized nitrogen in amides (GAFF2: n) and sp^3^-hybridized nitrogen (GAFF2: n3) are mostly separated along the second principal component, indicating a nuanced differentiation of nitrogen environments.

Similarly, for oxygen atoms, sp^2^-hybridized oxygen typically found in carbonyl groups (GAFF2: o) is largely separated for all models and molecular representations (Figure S4 in the Supplementary Information). However, in most configurations, the sp^2^-hybridized oxygen in hydroxyl groups (GAFF2: oh) and oxygen in ethers and esters (GAFF2: os) tend to overlap, except in the RoBERTa SMILES model, where these oxygen types are distinctly resolved.

We evaluated the impact of case sensitivity by modifying molecules to use only uppercase carbon atoms via the kekulize flag, thereby removing the distinction between aromatic (lowercase) and aliphatic (uppercase) carbons. As expected, this led to less distinct clustering of carbon atom types in the embedding space (Figure S7, Supplementary Information). Notably, despite the loss of case-based cues, the RoBERTa-based model still differentiated aromatic carbons (ca) from sp^2^-hybridized (c and c2) and sp^3^-hybridized (c3) carbons along the first principal component. In contrast, BART showed no such separation. This trend was consistent across other atom types as well: RoBERTa-based embeddings for nitrogen and oxygen atoms also displayed more structured separation than those from BART-based embeddings (Figure S8 and Figure S9 in the Supplementary Information).

We examined atom type clustering in embeddings produced by untrained models to better understand the extent to which structural information is captured without task-specific training. Using standard SMILES input, the untrained BART model already exhibited notable clustering across atom types (Figure S6 in the Supplementary Information). For carbon atoms, aromatic carbons (ca) and sp^3^-hybridized carbons (c3) were clearly separated along the first principal component, while sp^2^-hybridized aliphatic carbons (c2) and ketone/thioketone carbons (c) formed a more central, overlapping cluster. Similar, though less pronounced, trends were observed for nitrogen and oxygen atoms. This early clustering likely reflects inherent signals present in SMILES strings, such as lowercase characters denoting aromatic atoms and symbols like “=” indicating bond order, which can implicitly encode hybridization or bonding environments.

To test whether such early clustering persists when explicit aromaticity cues are removed, we repeated the analysis using kekulized SMILES input with the untrained BART model. In this setting, the case-based distinction between aromatic and aliphatic atoms is eliminated. Nonetheless, the model continued to show some chemically meaningful separation. For nitrogen atoms, sp^2^-hybridized atoms in amides (n) were distinguishable from sp^3^-hybridized atoms (n3) along the first principal component, while aromatic nitrogen (na) and nitrogen in amines (nh) remained more closely clustered. For oxygen atoms, sp^2^-hybridized oxygen (o) was clearly separated from oh and os (Figure S8, Figure S9 in the Supplementary Information).

These findings indicate that even without training, the structural information encoded in the input representations can lead to some initial clustering of atom types. This initial clustering is stronger when distinguishing between lowercase and uppercase c as is done in SMILES (Figure 5 and Figure S7 in the Supplementary Information). However, training significantly refines these embeddings, allowing models to develop a more nuanced understanding of atomic environments.

## 4 Discussion

In this study, we systematically investigated how three key design choices influence the performance and representational quality of large chemical language models. These choices include the choice of molecular representation (SMILES [1] vs. SELFIES [8]), the tokenisation strategy (atomwise [4] vs. SentencePiece [9]), and the underlying model architecture (BART [20] vs. RoBERTa [21]). By isolating and evaluating the effects of each of these variables across a range of tasks, we aimed to better understand how these architectural and preprocessing choices shape the ability of chemical language models to learn chemically meaningful representations.

Our comparative analysis was structured into four key components, each focusing on a different perspective of model evaluation. First, we evaluated the performance of each model configuration on downstream tasks after fine-tuning. Although certain configurations showed modest advantages, no configuration consistently outperformed the others across all tasks. This indicates that overall, the models learn robust internal representations that generalize well across different input representation and preprocessing strategies, highlighting the flexibility of these architectures in adapting to varied chemical representations. Second, we probed the latent space by training weaker classifiers to predict basic molecular properties using the embeddings generated by each model. The embedding of BART-based models consistently yielded better results than RoBERTa-based models, and SMILES-based models outperformed SELFIES-based models. The third analysis section once again inspected the latent space, but this time evaluating the UMAP and PCA plots of whole molecules. Our results indicate that the SMILES-based BART configuration achieved the best performance, with the molecular representation (SMILES) having a greater positive impact on performance than the specific model architecture. Finally, we examined how well the models captured atomic-level information by visualizing embeddings of atoms labelled by GAFF2 atom types. Here, RoBERTa showed a stronger ability to distinguish between atom types, and SMILES representations led to a more distinct separation than SELFIES, highlighting that the optimal configuration may vary depending on the granularity and nature of the chemical information being encoded.

Concerning preprocessing, we found that SentencePiece tokenization reduced training time by using shorter sequence lengths. However, this tokenization also sacrifices interpretability as its subword units do not correspond to tokens as easily interpretable as atoms from the atomwise tokenizer.

Overall, across our different analyses, SMILES representations consistently produced more chemically structured latent spaces compared to those derived from SELFIES. However, for certain generative tasks, where ensuring valid molecule generation is critical, the syntactic robustness of SELFIES may offer a distinct advantage [35]. The differences between BART and RoBERTa architecture were less pronounced in most analyses, where both yielded similarly interpretable latent spaces. Ultimately, the choice between BART and RoBERTa may be based on practical factors. RoBERTa is widely adopted, benefits from a more stable implementation, and has abundant community resources, making it a strong contender in many applications.

To further advance this research, future efforts could focus on several key areas. First, exploring the impact of alternative molecular representations, such as t-SMILES [36] and DeepSMILES [37], could lead to the development of more robust and generalizable models. Additionally, a direct comparison of BART’s generative capabilities with other large language model architectures, like GPT [38], would provide valuable insight into their respective strengths and limitations. Parallel to this, investigating the fundamental differences and potential synergies between large language models and graph-based approaches, such as GROVER [39], offers an exciting opportunity to expand the methodological toolkit for molecular modelling. Finally, utilizing highly curated datasets from specialized domains could reveal subtle chemical relationships and drive significant advancements in targeted applications.

## Supplementary Information

Supplementary File 1 provides extended figures and tables.

## Availability of data and materials

Pretraining data is taken from PubChem dataset [2] and fine-tuning dataset is taken from MoleculeNet [25, 26].

Implementation code is available on GitHub: https://github.com/ibmm-unibe-ch/SMILES_or_SELFIES.

## Acknowledgements

We thank Noah Kleinschmidt for careful review of the paper and valuable feedback on the project.

## Author contributions

**Inken Fender**: conceptualisation, data curation, formal analysis, investigation, methodology, software, validation, visualisation, writing **Jannik Adrian Gut**: conceptualisation, data curation, formal analysis, investigation, methodology, software, validation, visualisation, writing **Thomas Lemmin**: conceptualisation, investigation, funding acquisition, methodology, supervision, visualisation, writing

## References

[1] Weininger, D. Smiles, a chemical language and information system. Journal of chemical information and computer sciences 28, 31–36 (1988).

[2] Chithrananda, S., Grand, G. & Ramsundar, B. Chemberta: large-scale self-supervised pretraining for molecular property prediction. arXiv preprint arXiv:2010.09885 (2020).

[3] Ahmad, W., Simon, E., Chithrananda, S., Grand, G. & Ramsundar, B. Chemberta-2: Towards chemical foundation models. arXiv preprint arXiv:2209.01712 (2022).

[4] Schwaller, P. et al. Molecular Transformer: A Model for Uncertainty-Calibrated Chemical Reaction Prediction. ACS Central Science 5, 1572–1583 (2019).

[5] Chilingaryan, G. et al. Bartsmiles: Generative masked language models for molecular representations. Journal of Chemical Information and Modeling 64, 5832–5843 (2024).

[6] Sadeghi, S., Bui, A., Forooghi, A., Lu, J. & Ngom, A. Can large language models understand molecules? BMC Bioinformatics 25, 225 (2024).

[7] Ross, J. et al. Large-scale chemical language representations capture molecular structure and properties. Nature Machine Intelligence 4, 1256–1264 (2022). URL https://www.nature.com/articles/s42256-022-00580-7.

[8] Krenn, M., Hase, F., Nigam, A., Friederich, P. & Aspuru-Guzik, A. Self-referencing embedded strings (selfies): A 100% robust molecular string representation. Machine Learning: Science and Technology 1, 045024 (2020).

[9] Kudo, T. Sentencepiece: A simple and language independent subword tokenizer and detokenizer for neural text processing. arXiv preprint arXiv:1808.06226 (2018).

[10] Li, X. & Fourches, D. Smiles pair encoding: a data-driven substructure tokenization algorithm for deep learning. Journal of chemical information and modeling 61, 1560–1569 (2021).

[11] Krenn, M. et al. Selfies and the future of molecular string representations. Patterns 3 (2022).

[12] Leon, M., Perezhohin, Y., Peres, F., Popovic, A. & Castelli, M. Comparing smiles and selfies tokenization for enhanced chemical language modeling. Scientific Reports 14, 25016 (2024).

[13] Yuksel, A., Ulusoy, E., Unlu, A. & Dogan, T. Selformer: molecular representation learning via selfies language models. Machine Learning: Science and Technology 4, 025035 (2023).

[14] Flam-Shepherd, D., Zhu, K. & Aspuru-Guzik, A. Language models can learn complex molecular distributions. Nature Communications 13, 3293 (2022).

[15] Sultan, A., Sieg, J., Mathea, M. & Volkamer, A. Transformers for Molecular Property Prediction: Lessons Learned from the Past Five Years. Journal of Chemical Information and Modeling 64, 6259–6280 (2024).

[16] Kimber, T. B., Gagnebin, M. & Volkamer, A. Maxsmi: maximizing molecular property prediction performance with confidence estimation using smiles augmentation and deep learning. Artificial Intelligence in the Life Sciences 1, 100014 (2021).

[17] Kim, S. et al. Pubchem 2023 update. Nucleic acids research 51, D1373–D1380 (2023).

[18] Landrum, G. et al. rdkit/rdkit: 2024 09 5 (q3 2024) release (2025). URL 10.5281/zenodo.14779836.

[19] Wolf, T. et al. Huggingface’s transformers: State-of-the-art natural language processing. arXiv preprint arXiv:1910.03771 (2019).

[20] Lewis, M. et al. BART: Denoising sequence-to-sequence pre-training for natural language generation, translation, and comprehension. Proceedings of the 58th Annual Meeting of the Association for Computational Linguistics 7871–7880 (2020).

[21] Liu, Y. et al. Roberta: A robustly optimized bert pretraining approach. arXiv preprint arXiv:1907.11692 (2019).

[22] Vaswani, A. et al. Attention is all you need. Advances in neural information processing systems 30 (2017).

[23] Devlin, J., Chang, M.-W., Lee, K. & Toutanova, K. Bert: Pre-training of deep bidirectional transformers for language understanding. Proceedings of the 2019 conference of the North American chapter of the association for computational linguistics: human language technologies, volume 1 (long and short papers) 4171–4186 (2019).

[24] Ott, M. et al. fairseq: A fast, extensible toolkit for sequence modeling. arXiv preprint arXiv:1904.01038 (2019).

[25] Wu, Z. et al. Moleculenet: A benchmark for molecular machine learning. CoRR abs/1703.00564 (2017).

[26] Ramsundar, B. et al. Deep Learning for the Life Sciences (O’Reilly Media, 2019).

[27] Wang, J., Wang, W., Kollman, P. A. & Case, D. A. Automatic atom type and bond type perception in molecular mechanical calculations. Journal of Molecular Graphics and Modelling 25, 247–260 (2006).

[28] Kreyszig, E., Stroud, K. & Stephenson, G. Advanced engineering mathematics. Integration 9, 1014 (2008).

[29] Wilcoxon, F. Individual comparisons by ranking methods. Breakthroughs in statistics: Methodology and distribution 196–202 (1992).

[30] Pedregosa, F. et al. Scikit-learn: Machine learning in Python. Journal of Machine Learning Research 12, 2825–2830 (2011).

[31] Bishop, C. M. Neural networks for pattern recognition (Oxford university press, 1995).

[32] Cortes, C. & Vapnik, V. Support-vector networks. Machine learning 20, 273–297 (1995).

[33] Fan, R.-E., Chang, K.-W., Hsieh, C.-J., Wang, X.-R. & Lin, C.-J. Liblinear: A library for large linear classification. the Journal of machine Learning research 9, 1871–1874 (2008).

[34] McInnes, L., Healy, J. & Melville, J. Umap: Uniform manifold approximation and projection for dimension reduction. arXiv preprint arXiv:1802.03426 (2018).

[35] Shen, C., Krenn, M., Eppel, S. & Aspuru-Guzik, A. Deep molecular dreaming: inverse machine learning for de-novo molecular design and interpretability with surjective representations. Machine Learning: Science and Technology 2, 03LT02 (2021).

[36] Wu, J.-N. et al. t-smiles: a fragment-based molecular representation framework for de novo ligand design. Nature Communications 15, 4993 (2024).

[37] O’Boyle, N. & Dalke, A. Deepsmiles: An adaptation of smiles for use in machine-learning of chemical structures. chemrxiv preprint chemrxiv:7097960 (2018).

[38] Achiam, J. et al. Gpt-4 technical report. arXiv preprint arXiv:2303.08774 (2023).

[39] Rong, Y. et al. Self-supervised graph transformer on large-scale molecular data. Advances in neural information processing systems 33, 12559–12571 (2020).

[40] Loshchilov, I. & Hutter, F. Fixing weight decay regularization in adam. arXiv preprint arXiv:1711.05101 5, 5 (2017).

[41] Irwin, R., Dimitriadis, S., He, J. & Bjerrum, E. J. Chemformer: a pre-trained transformer for computational chemistry. Machine Learning: Science and Technology 3, 015022 (2022).

[42] Fabian, B. et al. Molecular representation learning with language models and domain-relevant auxiliary tasks. arXiv preprint arXiv:2011.13230 (2020).

