## Supplementary information for "Beyond performance: How design choices shape chemical language models"

| Argument | Value |
|---|---|
| batch-size | 32 |
| mask | 0.2 |
| tokens-per-sample | 512 |
| total-num-update | 500000 |
| warmup-updates | 1500 |
| task | denoising |
| arch | bart_base |
| optimizer | adam |
| lr-scheduler | polynomial_decay |
| lr | 1e-05 |
| dropout | 0.1 |
| criterion | cross_entropy |
| max-tokens | 3200 |
| weight-decay | 0.01 |
| attention-dropout | 0.2 |
| relu-dropout | 0.1 |
| share-decoder-input-output-embed | |
| share-all-embeddings | |
| clip-norm | 1.0 |
| attention-dropout | 0.2 |
| mask-length | span-poisson |
| replace-length | 1 |
| rotate | 0.0 |
| mask-random | 0.1 |
| permute-sentences | 1 |
| insert | 0 |
| poisson-lambda | 3.5 |
| GPU | 1xNVIDIA RTX A5000 |
| Training time | 5 days |

**Table S1** Fairseq pretraining parameters for BART models.

| Argument | Value |
|---|---|
| batch-size | 32 |
| tokens-per-sample | 512 |
| total-num-update | 500000 |
| warmup-updates | 1500 |
| task | masked_lm |
| arch | roberta_base |
| optimizer | adam |
| lr-scheduler | polynomial_decay |
| lr | 1e-05 |
| dropout | 0.1 |
| criterion | masked_lm |
| max-tokens | 3200 |
| weight-decay | 0.01 |
| attention-dropout | 0.2 |
| relu-dropout | 0.1 |
| clip-norm | 1.0 |
| attention-dropout | 0.2 |
| GPU | 1xNVIDIA RTX A5000 |
| Training time | 4 days |

**Table S2** Fairseq pretraining parameters for RoBERTa models.

### Supplementary Information S1 Training details

The BART and and RoBERTa pretraining was conducted in accordance with the fairseq [1] parameters specified in Table S1 and Table S2 respectively. Further pre-training information can be found in our public Weights and Biases workspace https://wandb.ai/ibmm-lemmin/pre-train/overview.

Fine-tuning was conducted with the AdamW[2] optimiser, utilising Adam-betas (0.9, 0.999), Adam-eps 1e-08, weight decay 0.01, dropout 0.2 and clip norm 0.1. In order to identify the optimal learning rates, a range of [1e-05, 5e-05, 5e-06] was tested with one seed each time. In the end, five seeds were used with the best learning rate to assess the performance and gather more data points for the z-score. Further information can be found on Weights and Biases: pretrain, tox21, lipo, hiv, delaney, clintox, clearance, bbbp, bace regression, bace classification.

| Representation | Tokenizer | Chirality | Language model | BACE ↑ | BBBP ↑ | Clintox ↑ | HIV ↑ | Tox21 ↑ |
|---|---|---|---|---|---|---|---|---|
| SELFIES | Atom | Explicit | BART | 0.769±0.011 | 0.701±0.032 | 0.730±0.099 | 0.753±0.022 | 0.671±0.028 |
| | | | RoBERTa | 0.804±0.019 | 0.701±0.010 | **0.880±0.031** | 0.766±0.014 | 0.639±0.044 |
| | | Implicit | BART | 0.784±0.016 | 0.685±0.018 | 0.608±0.031 | 0.754±0.010 | 0.695±0.016 |
| | | | RoBERTa | 0.763±0.088 | 0.705±0.017 | 0.876±0.008 | 0.730±0.019 | 0.638±0.043 |
| | SentencePiece | Explicit | BART | 0.793±0.018 | 0.698±0.014 | 0.657±0.053 | 0.724±0.010 | **0.719±0.013** |
| | | | RoBERTa | 0.806±0.021 | 0.702±0.010 | 0.788±0.045 | 0.733±0.014 | 0.703±0.018 |
| | | Implicit | BART | 0.684±0.021 | 0.704±0.018 | 0.611±0.032 | 0.740±0.006 | 0.673±0.018 |
| | | | RoBERTa | 0.675±0.097 | 0.691±0.009 | 0.689±0.030 | 0.716±0.023 | 0.672±0.007 |
| SMILES | Atom | Explicit | BART | **0.824±0.026** | 0.695±0.028 | 0.743±0.043 | 0.763±0.013 | 0.659±0.038 |
| | | | RoBERTa | 0.701±0.056 | **0.746±0.012** | 0.844±0.038 | 0.753±0.016 | 0.640±0.018 |
| | | Implicit | BART | 0.798±0.016 | 0.692±0.020 | 0.730±0.112 | 0.757±0.013 | 0.687±0.016 |
| | | | RoBERTa | 0.736±0.042 | 0.727±0.021 | 0.726±0.103 | 0.754±0.004 | 0.674±0.012 |
| | SentencePiece | Explicit | BART | 0.788±0.028 | 0.685±0.009 | 0.611±0.038 | **0.778±0.011** | 0.698±0.035 |
| | | | RoBERTa | 0.776±0.009 | 0.729±0.009 | 0.644±0.018 | 0.771±0.010 | 0.667±0.057 |
| | | Implicit | BART | 0.717±0.098 | 0.714±0.005 | 0.593±0.071 | 0.759±0.023 | 0.696±0.017 |
| | | | RoBERTa | 0.612±0.046 | 0.697±0.013 | 0.666±0.046 | 0.750±0.007 | 0.698±0.010 |
| ChemBERTa-2 [3] | | | | 0.799 | 0.742 | 0.601 | | 0.834 |

**Table S3** Downstream classification ROC AUC score mean and standard deviation scores of five seeds on MoleculeNet datasets [4] across different molecular representations, tokenizers, chirality representations and language models

| Representation | Tokenizer | Chirality | Language model | BACE ↓ | Clearance ↓ | Delaney ↓ | Lipo ↓ |
|---|---|---|---|---|---|---|---|
| SELFIES | Atom | Explicit | BART | 0.878±0.144 | 1.159±0.033 | 0.578±0.017 | 0.701±0.019 |
| | | | RoBERTa | 1.159±0.145 | 1.241±0.102 | 0.480±0.030 | 0.731±0.022 |
| | | Implicit | BART | 0.853±0.180 | 1.182±0.023 | 0.608±0.036 | 0.678±0.016 |
| | | | RoBERTa | 1.061±0.234 | 1.200±0.019 | **0.454±0.017** | 0.693±0.018 |
| | SentencePiece | Explicit | BART | 1.161±0.101 | 1.175±0.026 | 0.507±0.026 | 0.671±0.012 |
| | | | RoBERTa | 0.902±0.175 | 1.206±0.059 | 0.513±0.015 | 0.734±0.010 |
| | | Implicit | BART | 1.039±0.170 | 1.243±0.039 | 0.529±0.022 | 0.705±0.007 |
| | | | RoBERTa | 0.892±0.052 | 1.242±0.053 | 0.513±0.025 | 0.751±0.040 |
| SMILES | Atom | Explicit | BART | **0.842±0.072** | 1.145±0.026 | 0.520±0.053 | 0.672±0.012 |
| | | | RoBERTa | 1.085±0.159 | 1.209±0.080 | 0.458±0.020 | 0.673±0.007 |
| | | Implicit | BART | 0.854±0.184 | 1.182±0.015 | 0.457±0.021 | 0.677±0.021 |
| | | | RoBERTa | 0.995±0.127 | 1.171±0.030 | 0.468±0.022 | **0.653±0.015** |
| | SentencePiece | Explicit | BART | 0.888±0.193 | 1.172±0.032 | 0.550±0.029 | 0.689±0.011 |
| | | | RoBERTa | 0.995±0.092 | 1.229±0.041 | 0.518±0.007 | 0.722±0.015 |
| | | Implicit | BART | 0.978±0.164 | **1.134±0.043** | 0.556±0.047 | 0.698±0.012 |
| | | | RoBERTa | 1.020±0.099 | 1.179±0.076 | 0.508±0.019 | 0.712±0.022 |
| MolBERT [5] | | | | | 0.948 | 0.531 | 0.561 |
| ChemFormer [6] | | | | | 1.230 | 0.633 | 0.598 |

**Table S4** Downstream regression RMSE score mean and standard deviation from five seeds on MoleculeNet datasets [4] across different molecular representations, tokenizers, chirality representations and language models

| Better choice | Worse choice | p-value |
|---|---|---|
| SMILES | SELFIES | 0.004 |
| Atomwise | SentencePiece | 0.020 |
| BART | RoBERTa | 0.416 |
| Explicit | Implicit | 0.123 |

**Table S5** One-sided Wilcoxon signed-rank test [7] using matching mean z-scores of all seven downstream tasks and all 16 configurations each, only differing in the indicated parameter of interest. This means for each test, we had two matching cohorts of 56 measurements. The first column indicates the better performing choice compared to the second column, while the third indicates the p-value of the hypothesis that the worse choice performs better than the better choice. Comparatively, 1-(p-value) is the p-value for the hypothesis that the better choice outperforms the worse choice. Regression scores have been multiplied by -1 to have the same ranking direction.

### Supplementary Information S2 Simpler classifier details

To train the weak classifiers we used the implementations from scikit-learn [8]. For k-nearest-neighbours classifier we searched the hyperparameters for n_neighbours [1, 5, 11] and weights [uniform, distance] and for the SVC and LinearSVC we used a C in [0.1, 1, 10], additionally we used LinearSVC with max_iter=1000. The splitting into train and test set was done randomly.

### References


[1] Ott, M. *et al.* fairseq: A fast, extensible toolkit for sequence modeling. *arXiv preprint arXiv:1904.01038* (2019).

[2] Loshchilov, I. & Hutter, F. Fixing weight decay regularization in adam. *arXiv preprint arXiv:1711.05101* **5**, 5 (2017).

[3] Ahmad, W., Simon, E., Chithrananda, S., Grand, G. & Ramsundar, B. Chemberta-2: Towards chemical foundation models. *arXiv preprint arXiv:2209.01712* (2022).

[4] Wu, Z. *et al.* Moleculenet: A benchmark for molecular machine learning. *CoRR* **abs/1703.00564** (2017).

[5] Fabian, B. *et al.* Molecular representation learning with language models and domain-relevant auxiliary tasks. *arXiv preprint arXiv:2011.13230* (2020).

[6] Irwin, R., Dimitriadis, S., He, J. & Bjerrum, E. J. Chemformer: a pre-trained transformer for computational chemistry. *Machine Learning: Science and Technology*

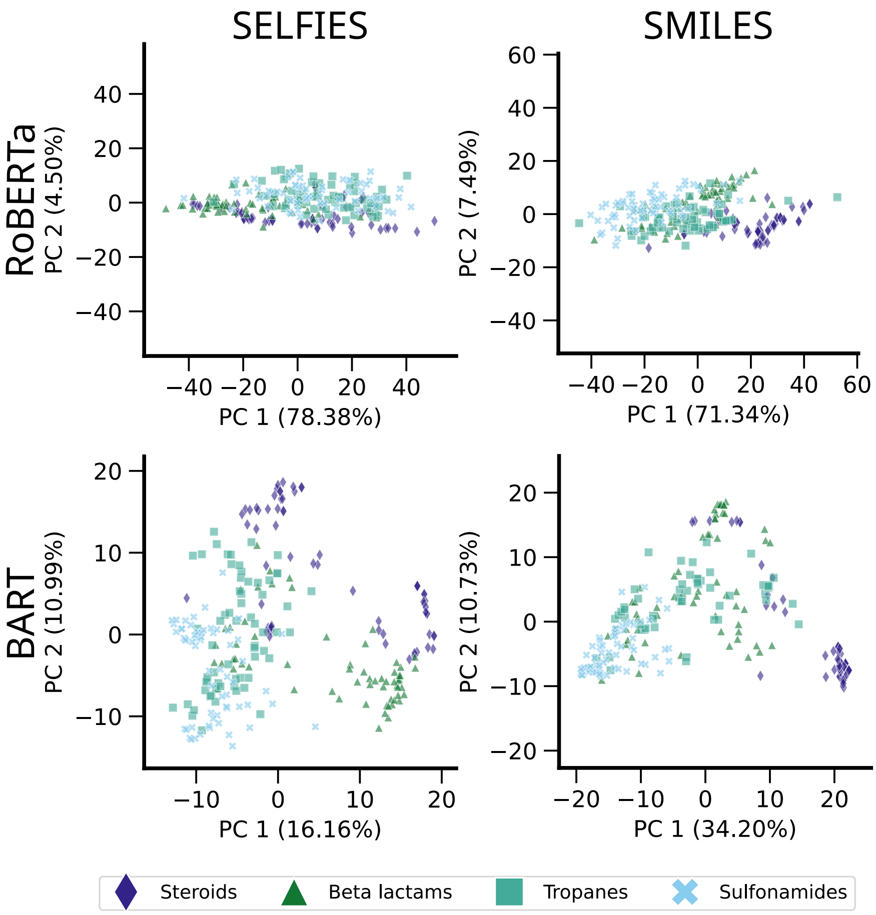


**Fig. S1** PCA embeddings of various molecules using SELFIES or SMILES, RoBERTa or BART, atomwise tokeniser and implicit isomer representation

**3**, 015022 (2022).

[7] Wilcoxon, F. in *Individual comparisons by ranking methods* 196–202 (Springer, 1992).

[8] Pedregosa, F. *et al.* Scikit-learn: Machine learning in Python. *Journal of Machine Learning Research* **12**, 2825–2830 (2011).

[9] Wang, J., Wang, W., Kollman, P. A. & Case, D. A. Automatic atom type and bond type perception in molecular mechanical calculations. *Journal of Molecular Graphics and Modelling* **25**, 247–260 (2006).

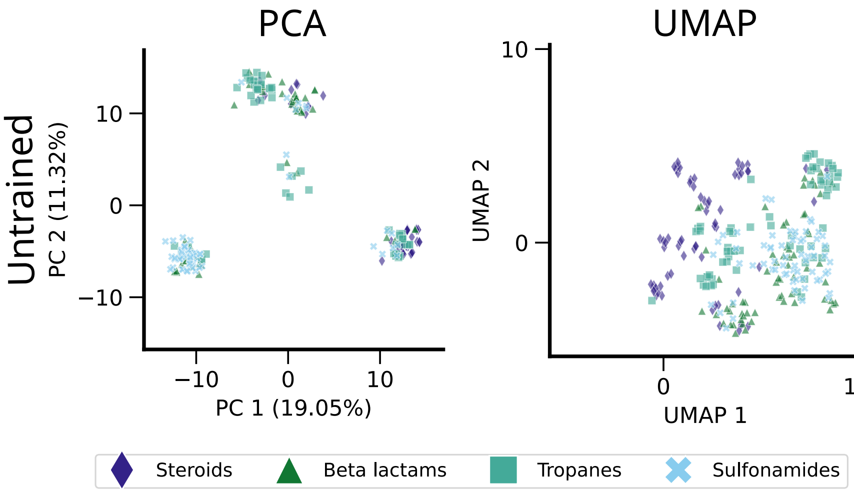


**Fig. S2** PCA and UMAP embeddings of various molecules using an untrained SMILES model

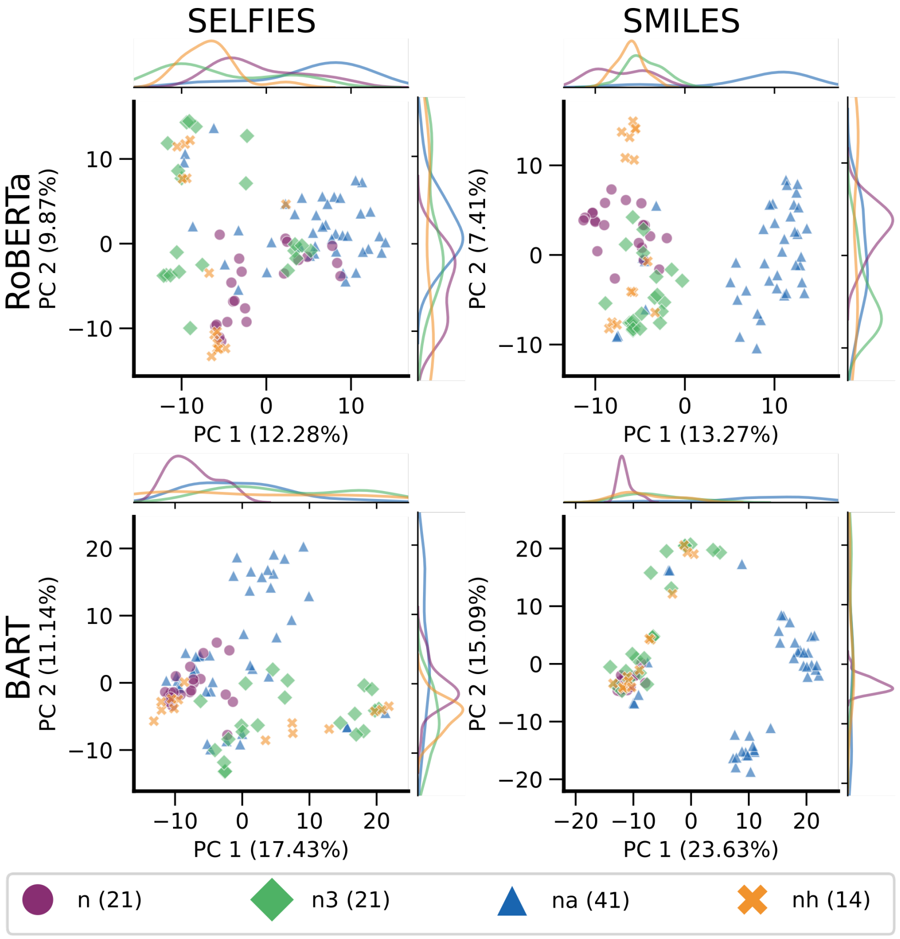


**Fig. S3** PCA of nitrogen atom type embeddings of SMILES or SELFIES-based models BART and RoBERTA with atomwise tokeniser and implicit chirality. The GAFF2 atom types have been determined by antechamber[9] and correspond to the following hybridizations: n: $sp^2$ in amide, n3: $sp^3$ N with 3 substitutions, na: $sp^2$ N with 3 substitutions, nh: amine N connected to the aromatic rings. Amount of samples in brackets.

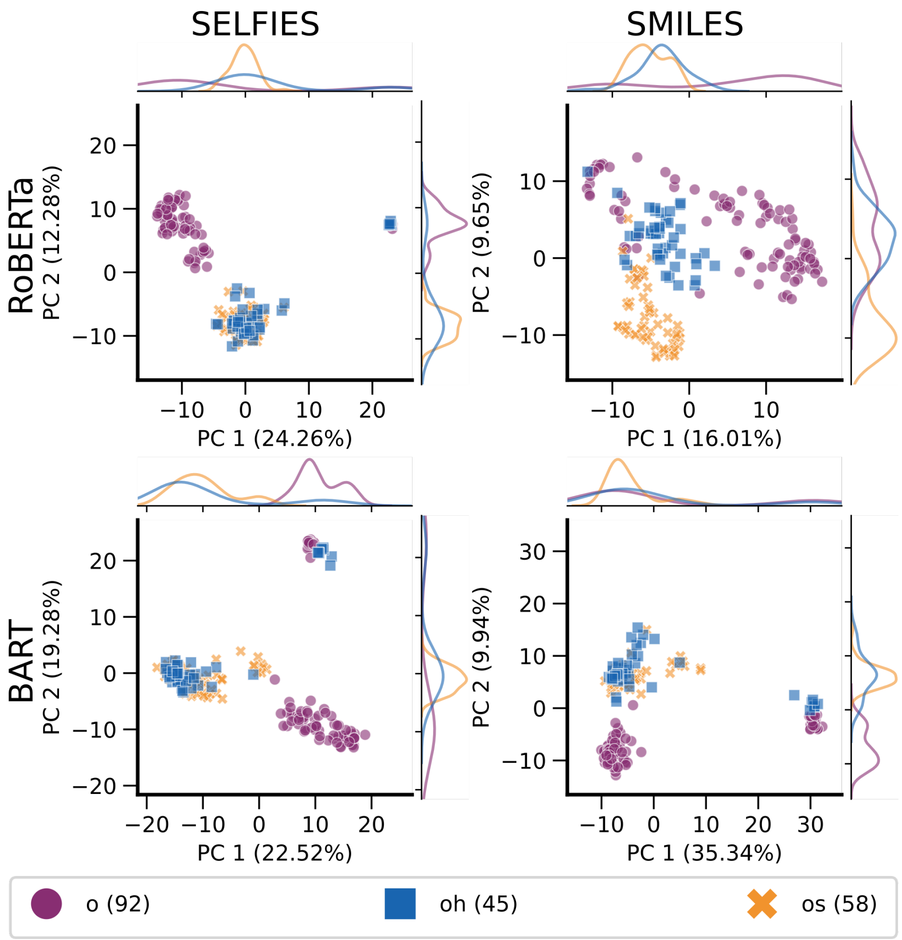


**Fig. S4** PCA of oxygen atom type embeddings of SMILES or SELFIES-based models BART and RoBERTA with atomwise tokeniser and implicit chirality. The GAFF2 atom types have been determined by antechamber[9] and correspond to the following hybridizations: o: $sp^2$ O in C=O and COO-, oh: $sp^3$ O in hydroxyl group, os: $sp^3$ O in ether and ester. Amount of samples in brackets.

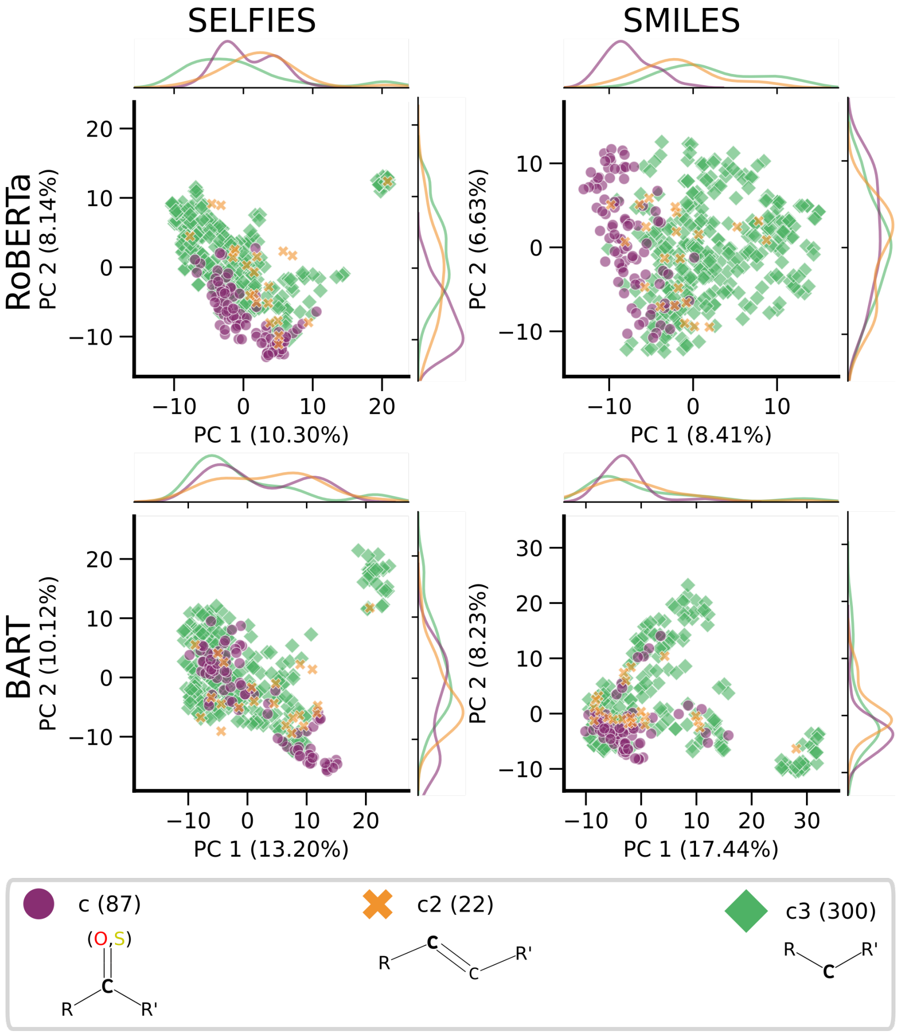


**Fig. S5** PCA of non-aromatic carbon atom type embeddings of SMILES or SELFIES-based models BART and RoBERTA with atomwise tokeniser and implicit chirality. The GAFF2 atom types have been determined by antechamber[9] and correspond to the following hybridizations: c: $sp^2$ in C=O, C=S, c2: $sp^2$ in aliphatic carbon, c3: $sp^3$. Amount of samples in brackets.

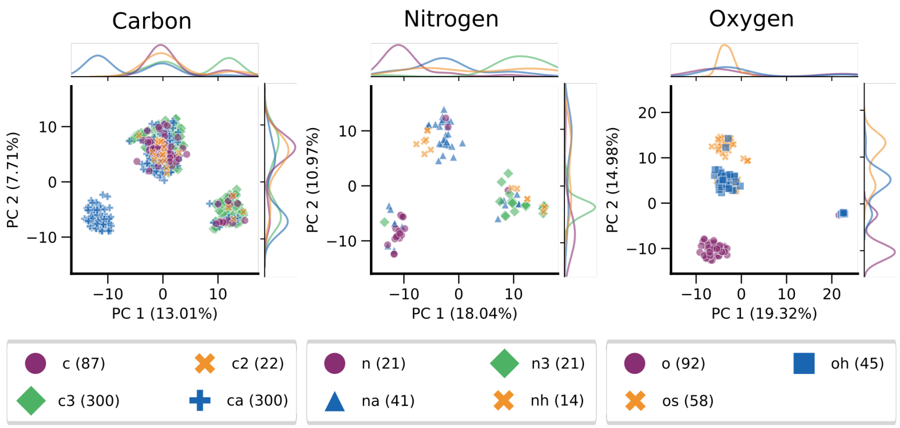


**Fig. S6** PCA of embeddings of carbon, nitrogen, and oxygen atom types from untrained SMILES-BART with atomwise tokeniser and implicit chirality. The GAFF2 atom types have been determined by antechamber[9] and correspond to the following hybridizations: c: $sp^2$ in C=O, C=S, c2: $sp^2$ in aliphatic carbon, c3: $sp^3$, n: $sp^2$ in amide, n3: $sp^3$ N with 3 substitutions, na: $sp^2$ N with 3 substitutions, nh: amine N connected to the aromatic rings, o: $sp^2$ O in C=O and COO-, oh: $sp^3$ O in hydroxyl group, os: $sp^3$ O in ether and ester. Amount of samples in brackets.

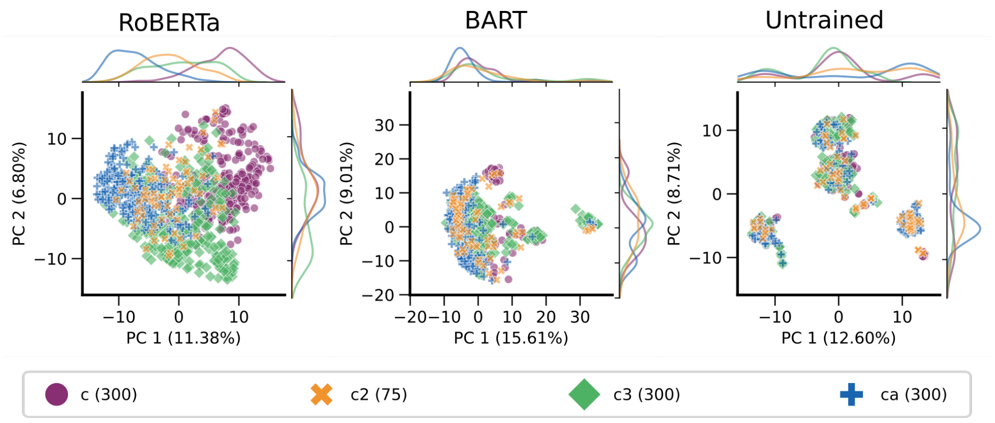


**Fig. S7** PCA of carbon atom type embeddings of kekulized SMILES that contain only uppercase carbons from models BART and RoBERTA with atomwise tokeniser and implicit chirality and the untrained BART. The GAFF2 atom types have been determined by antechamber[9] and correspond to the following hybridizations: c: $sp^2$ in C=O, C=S, c2: $sp^2$ in aliphatic carbon, c3: $sp^3$, ca: $sp^2$ in aromatic carbon. Amount of samples in brackets (Note: Since mapping to SELFIES was not done here, more atom types could be analysed and so numbers of atom types differ to previous plots.)

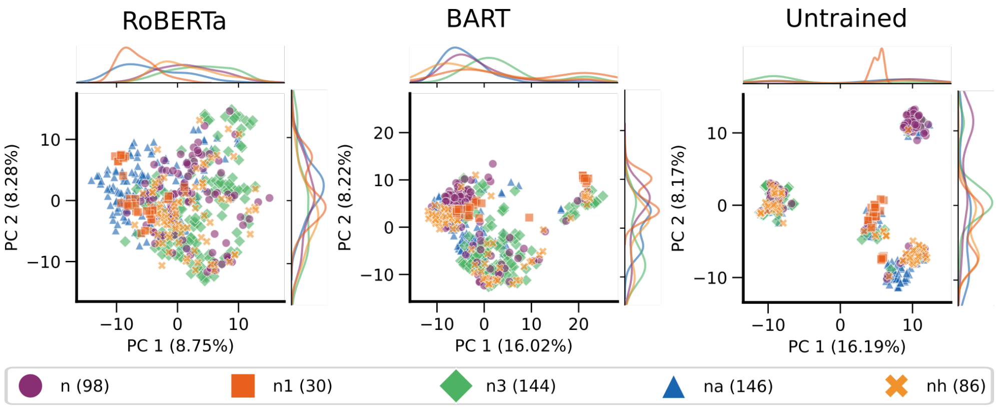


**Fig. S8** PCA of nitrogen atom type embeddings of kekulized SMILES that contain only uppercase carbons from models BART and RoBERTA with atomwise tokeniser and implicit chirality and the untrained BART. The GAFF2 atom types have been determined by antechamber[9] and correspond to the following hybridizations: n: $sp^2$ in amide, n1: $sp^1$ N, n3: $sp^3$ N with 3 substitutions, na: $sp^2$ N with 3 substitutions, nh: amine N connected to the aromatic rings. Amount of samples in brackets (Note: Since mapping to SELFIES was not done here, more atom types could be analysed and so numbers of atom types differ to previous plots.)

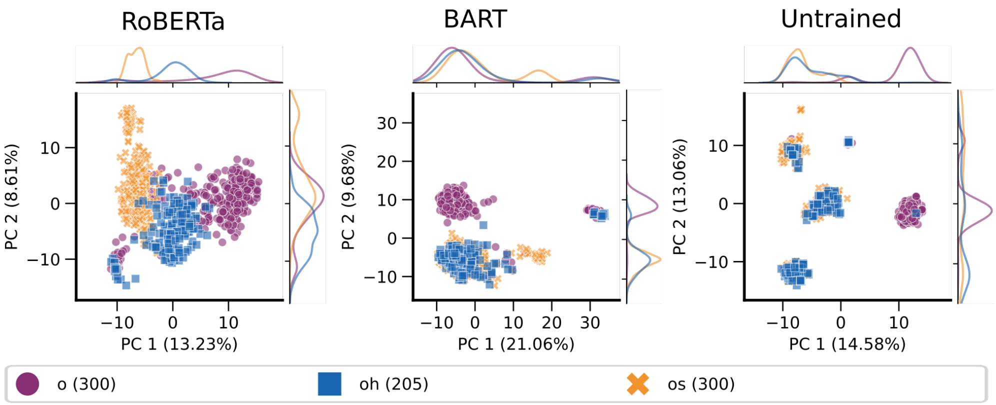


**Fig. S9** PCA of oxygen atom type embeddings of kekulized SMILES that contain only uppercase carbons from models BART and RoBERTA with atomwise tokeniser and implicit chirality and the untrained BART. The GAFF2 atom types have been determined by antechamber[9] and correspond to the following hybridizations: o: $sp^2$ O in C=O and COO-, oh: $sp^3$ O in hydroxyl group, os: $sp^3$ O in ether and ester.. Amount of samples in brackets (Note: Since mapping to SELFIES was not done here, more atom types could be analysed and so numbers of atom types differ to previous plots.)
